# The Burger Battle: The Future is Blended

**DOI:** 10.64898/2026.06.17.732995

**Authors:** Aeneas O Koosis, Christopher D Gardner, Ellen Kuhl

**Author notes:** Corresponding author: *Email address:* (Aeneas O Koosis).

## Abstract

Texture is one of the major barriers to acceptance of sustainable protein products, yet the relationship between instrumental texture and consumer perception remains poorly understood. Here we combined consumer sensory evaluation and texture profile analysis to identify the factors that drive burger acceptance across blended, animal-based, and plant-based formulations. A total of 472 diners evaluated beef-mushroom (*n* = 171), turkey (*n* = 100), bean (*n* = 100), and pea (*n* = 101) burgers served in Stanford University dining halls, while texture profile analysis quantified physical texture (*n* = 10 per product). The beef-mushroom burger achieved the highest ratings for meatiness (77.0/100), tastiness (69.4/100), and moistness (61.8/100), and 78% of consumers rated its moistness as just-about-right. Although the pea burger exhibited mechanical properties similar to the beef-mushroom burger, including indistinguishable cohesiveness values (0.53 vs. 0.53), it received substantially lower ratings for meatiness (56.1 vs. 77.0), indicating that mechanical similarity alone does not ensure consumer acceptance. Across products, moistness emerged as the primary sensory limitation, with 58% of turkey consumers and 46% of bean consumers reporting insufficient moisture. Cohesiveness showed the strongest association with perceived meatiness (*r* = 0.82). These findings demonstrate that successful burger reformulation requires more than mechanical matching; products must reproduce the moisture, flavor, and eating experience associated with meat. Blended burgers may represent the most practical near-term strategy for reducing meat consumption without compromising eating quality.

## 1. Motivation

The global food system must reduce its environmental footprint while continuing to provide appealing and nutritionally adequate sources of dietary protein. Conventional beef production contributes substantially to greenhouse gas emissions, land use, and water consumption [1]. Plant-based and blended meat products could reduce the environmental impact of protein production while increasing dietary fiber and lowering saturated fat intake [2]. Yet, widespread adoption of these products remains limited because consumers often perceive them as inferior to animal-derived meat in texture and eating experience [3].

Texture represents one of the primary barriers to consumer acceptance of meat alternatives [4]. While environmental and health concerns may motivate initial product trial, repeat purchase behavior depends predominantly on sensory satisfaction [5]. Consumers evaluate meat alternatives through attributes such as juiciness, chewiness, softness, and fibrousness [6]. These attributes strongly influence whether a product feels meat-like and acceptable. Texture parity may therefore represent a necessary condition for broad consumer adoption of sustainable protein products [7].

Burgers represent a uniquely high-leverage intervention point in the American diet [8]. On any given day, 21.4% of the U.S. population consumes a beef sandwich, and hamburgers and cheeseburgers account for 17.3% of all unprocessed red meat intake [9]. Beef burgers contribute 6.3% of mean daily energy intake across the U.S. population and 26.2% on days of consumption [10]. These products also contribute approximately 10% of Americans’ saturated fat and sodium intake [10]. Compared to plant-based alternatives, conventional beef burgers contain more calories, saturated fat, and cholesterol, while providing less fiber [11]. This nutritional profile raises concern because most Americans fail to meet dietary fiber recommendations and overconsume saturated fat and cholesterol [12]. These patterns suggest that even modest reformulation of burger products could yield substantial nutritional and environmental benefits at population scale [13].

Blended burgers that partially replace meat with plant-based ingredients such as mushrooms may provide a practical transition strategy for consumers unwilling to adopt fully plant-based alternatives [14]. By retaining animal-derived texture and flavor cues while reducing meat content, blended formulations may achieve greater consumer acceptance than fully plant-based products [15]. Recent studies suggest that blended products can match or exceed conventional meat products in blind sensory evaluations [16]. Consumers also express greater purchase intent for blended products than for fully plant-based alternatives [17].

Despite advances in food science and processing technology, few studies have examined consumer sensory responses to diverse burger formulations, including beef-mushroom blends, poultry-based, plant-based, and vegetable-based options, in direct comparison using both subjective consumer evaluation and objective instrumental texture analysis [15]. Recent studies on balanced protein products suggests that blending animal and plant proteins can achieve taste parity with conventional meat [16], and that consumers are 40% more likely to express purchase interest in blended products compared to fully plant-based alternatives [17]. However, the specific textural attributes driving acceptance remain under examined. Product developers seeking to optimize formulations need to understand these attributes [18]. Correlating consumer perception with instrumental texture profile analysis parameters provides mechanistic insights into the physical basis of sensory experience [19].

*Here we test the hypothesis that specific mechanical texture properties, particularly cohesiveness and moisture retention, govern consumer perception of meatiness and overall acceptance across plant-based, blended, and animal-based burgers*.

To address this question, we combine consumer sensory evaluation with rheological texture profile analysis across four burger products: beef-mushroom, turkey, bean, and pea protein. We first quantify consumer perception of key sensory attributes including meatiness, moistness, chewiness, softness, and tastiness. We then characterize the mechanical response of each burger through instrumental texture profile analysis and correlate these measurements with consumer perception. Together, these analyses identify the textural characteristics that most strongly predict consumer acceptance and provide mechanistic targets for future burger formulation.

## 2. Materials and Methods

### 2.1. Participants

We recruited participants from the Stanford University community during March–April 2026. Recruitment materials invited adults to participate in a “burger taste test study” in the dining hall setting. Inclusion criteria required participants to be 18 years or older, able to read and understand English, and willing to consume beef, turkey, and plant-based burger products. Exclusion criteria included known food allergies to study ingredients including soy, wheat, pea protein, mushrooms, and other relevant allergens, and self-reported taste or smell disorders. A target sample size of *n* = 400 participants provided adequate statistical power (1 - *β* = 0.80) to detect medium effect sizes (Cohen’s d = 0.5) in texture perception differences across product conditions at *α* = 0.05. We ultimately enrolled 472 participants who completed all study procedures. Participants provided electronic informed consent through Qualtrics and received a $10 gift card after survey completion. The Stanford University Institutional Review Board approved the study (eProtocol #84659) as exempt under taste and food quality evaluation (45 CFR 46.104(d)(6)).

### 2.2. Burger Products

We evaluated four burger products that represented diverse protein sources and formulation strategies (Table 1).

**Table 1:** Burger products evaluated in this study. We selected products based on availability within the Stanford dining system. The beef-mushroom blend contained approximately 90% grass-fed beef and 10% cremini mushrooms by weight. The university food service provider supplied the turkey, bean, and pea protein patties as commercial products. Dining hall staff labeled all products by name at the serving station, and participants remained aware of product identity during selection.

| Product | Label | Protein Source | Key Ingredients |
| --- | --- | --- | --- |
| beef-mushroom | beef | Grass-fed beef + mushroom | Grass-fed beef, cremini mushrooms, onions, spices |
| turkey | turkey | Poultry | Turkey, rosemary, sage |
| bean | bean | Legume/vegetable | Black beans, corn, oats,peppers, onions, spices |
| pea | pea | Plant protein isolates | Pea protein, brown rice protein, coconut oil, beet juice |

### 2.3. Sample Preparation

Dining hall staff cooked all burger patties (approximately 110–140 g each) on a commercial flat-top grill to an internal temperature of approximately 71^°^C (160^°^F) for beef-mushroom and turkey patties and 74^°^C (165^°^F) for plant-based patties. Staff held cooked patties in steam-table trays at approximately 60^°^C (140^°^F) for up to 30 minutes before replacement. The dining halls used a build-your-own burger format in which participants selected their preferred patty from labeled serving trays (Figure 1).

**Figure 1:**
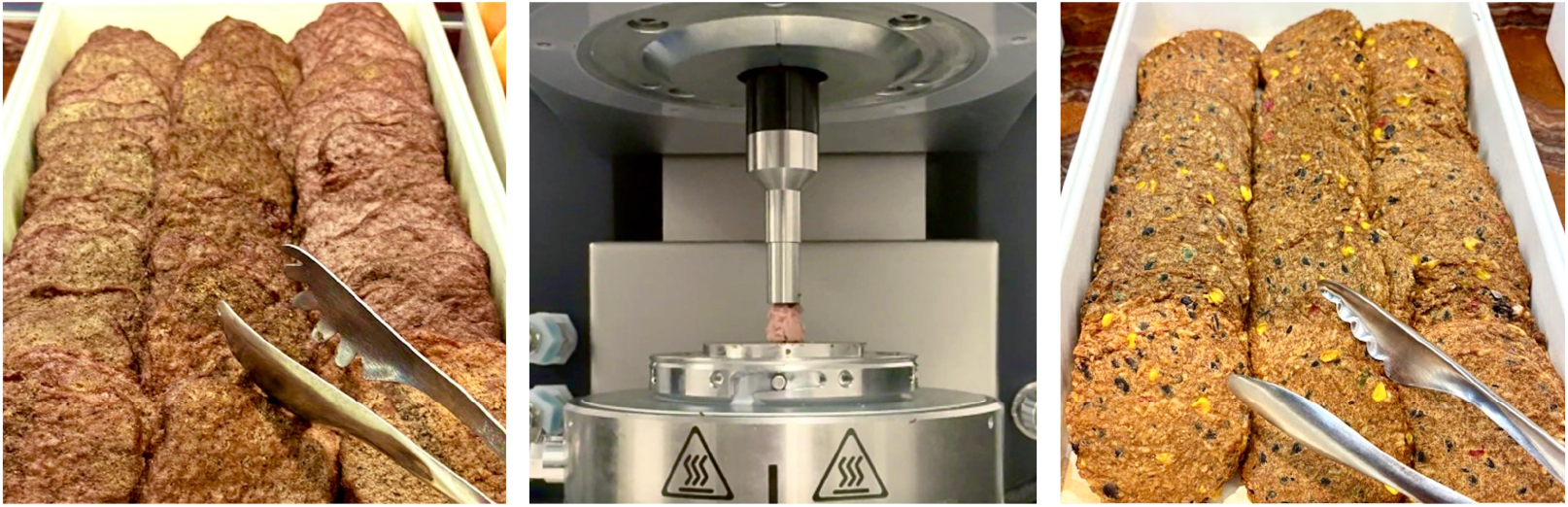
Study setup and texture profile analysis. Standardized burger presentation during consumer sensory evaluation in Stanford dining halls, including beef-mushroom (left) and bean burger (right). Participants select burgers under normal dining conditions and complete sensory surveys on personal devices. We complement these surveys with a texture profile analysis (middle). The experimental setup shows a cylindrical burger sample mounted between the compression plates of a rheometer for double-compression testing.

### 2.4. Consumer Sensory Evaluation

#### 2.4.1. Procedure

We conducted sensory evaluations in multiple campus dining halls during regular meal service hours. Participants first selected a burger as part of their normal meal choice and then completed a voluntary survey about the selected product. Participants remained seated at standard dining hall tables and accessed the study information sheet and survey through a QR code or web link. After providing informed consent, participants consumed the selected burger and completed the survey on personal devices. Participants assembled their own burgers and could add toppings and condiments according to personal preference, consistent with standard dining hall practice. We adopted a between-subjects design in which each participant evaluated only one burger type to reflect naturalistic dining behavior in dining halls, where consumers typically select a single entrée and customize it before consumption.

#### 2.4.2. Sensory Attribute Ratings

Participants rated each burger sample on eight texture and flavor attributes using visual analog scales ranging from 0 (lowest intensity) to 100 (highest intensity). The evaluated attributes included *moist, chewy, fatty, fibrous, hard, soft, tasty*, and *meaty* [20]. Participants also rated the burger overall on a 7-point scale ranging from *very poor* to *exceptional*.

#### 2.4.3. Just-About-Right (JAR)

Participants rated five burger attributes, including *moist, chewy, fatty, fibrous*, and *savory*, using a 3-point Just-About-Right scale (1 = not enough, 2 = just about right, 3 = too much). We selected the 3-point format because penalty analysis collapses responses into three functional categories regardless of original scale length [21]. The simplified scale reduced respondent burden while preserving diagnostic power for identifying reformulation priorities [22].

#### 2.4.4. Meat Attachment Questionnaire

We assessed psychological attachment to meat using the 5-item dependence subscale of the Meat Attachment Questionnaire (MAQ) [23]. Participants responded to five statements: (i) I do not picture myself without eating meat regularly; (ii) If I could not eat meat I would feel weak; (iii) I would feel fine with a meatless diet [reverse-scored]; (iv) If I was forced to stop eating meat I would feel weak; and (v) Meat is irreplaceable in my diet. Participants rated each item on a 5-point Likert scale ranging from strongly 1 (disagree) to 5 (strongly agree). We calculated the composite MAQ score as the mean across all five items.

#### 2.4.5. Choice Factor Importance

Participants rated the importance of eight factors in burger selection using visual analog scales ranging from 0 to 100. The evaluated factors included *taste, texture, price, health, environmental impact, animal welfare, convenience*, and *familiarity*.

#### 2.4.6. Facilitators of Plant-Based Meat Selection

To identify interventions that could increase plant-based meat consumption, participants completed a randomized check-all-that-apply (CATA) question that evaluated six factors: *price parity, sensory equivalence, default menu placement, nutritional equivalence, social normalization*, and *reduced use of highly processed additives*. Participants could also provide additional responses through an open-ended *other* option.

### 2.5. Texture Profile Analysis

We performed texture profile analysis using a TA Instruments Discovery HR-2 rheometer equipped with an 8 mm diameter flat compression plate. We excised standardized cylindrical samples with 8 mm diameter from the center of cooked burger patties and maintained samples at room temperature (22^°^C) before testing. The testing protocol included two consecutive compression cycles to 50% strain with a recovery interval between compressions [24]. The rheometer recorded force-displacement data at a sampling frequency of 244 Hz. We processed raw force data using Gaussian filtering (σ = 2) to reduce noise while preserving peak characteristics. We identified compression cycle boundaries using a 0.1 N force threshold [15]. We extracted six texture profile analysis parameters from the force-time curves [19]: *stiffness* as the initial slope of the force-displacement curve, measured in in kiloPascal, calculated as *F*_1_/(ε*A*) where *F* is peak force, *A* is sample area, and ε is strain; *hardness* as the maximum force during the first compression cycle *F*, measured in Newton; *cohesiveness* as the ratio of the positive force area during second compression to that during the first, (*A*_3_ + *A*_4_)/(*A*_1_ + *A*_2_); *springiness* as the ratio of time to achieve peak force during second compression to that during the first, *t*_2_/*t*_1_; *resilience* as the ratio of energy recovered during unloading to the energy expended during loading in the first cycle, *A*_2_/*A*_1_; and *chewiness* as the product of hardness, cohesiveness, and springiness, measured in Newton. We performed at least 10 replicate measurements for each burger type (*n* = 40 total samples).

### 2.6. Statistical Analysis

We conducted all statistical analyses in R version 4.3.2. We calculated descriptive statistics, including means and standard deviations, for all sensory attributes and texture profile analysis parameters. We used Kruskal-Wallis H-tests to compare sensory ratings across the four burger conditions and performed post-hoc pairwise comparisons using Mann-Whitney U tests with Bonferroni correction. We selected the Kruskal-Wallis test because sensory ratings followed ordinal scales and did not require assumptions of normality. For texture profile analysis parameters, we used one-way ANOVA with Tukey’s honestly significant difference (HSD) tests because the instrumental measurements were continuous variables. We performed penalty analysis to quantify how deviations from just about right influenced overall liking. We defined the penalty as the difference in mean overall liking between participants who rated an attribute as *just about right* and those who rated the same attribute as *not enough* or *too much*. Larger penalties identified priority targets for product reformulation. We defined actionable deviations as penalties ≥0.50 on the 7-point overall liking scale with *p* < 0.05, following standard industry thresholds [21]. We calculated Pearson correlation coefficients (*r*) to examine relationships between instrumental texture profile analysis parameters and consumer sensory ratings at the burger-product level (*n* = 4 burger types). We conducted mean-drop analysis according to established industry protocols [21]. We flagged penalties ≥0.50 on the 7-point overall liking scale as actionable. We defined statistical significance as *p* < 0.05 for all analyses.

## 3. Results

### 3.1. Participant Characteristics

Table 2 summarizes the demographics the *n* = 472 participants who completed the study. Participants selected all four burgers in approximately balanced proportions, with the beef-mushroom burger representing the most frequently selected option (36.2%). The cohort consisted predominantly of young adults between 18 and 24 years of age (78.0%) and identified primarily as omnivores (85.6%). Mean meat attachment questionnaire (MAQ) scores indicated moderate psychological attachment to meat consumption across the study population (3.31 ± 0.91), while participants reported that plant-based foods comprised 36.1 ± 22.4% of their diets.

**Table 2:** Participant demographics and dietary characteristics. Distribution of burger selection, age, gender, ethnicity, dietary pattern, and meat attachment questionnaire (MAQ) scores across the study cohort (*N* = 472). Values represent counts with percentages or mean ± standard deviation. Participants predominantly identified as omnivores and selected all four burger types in approximately balanced proportions.

| Characteristic | $n$ (%) | | $n$ (%) |
| --- | --- | --- | --- |
| **Burger Type** |  |  |  |
| beef-mushroom | 171 (36.2%) | bean | 100 (21.2%) |
| turkey | 100 (21.2%) | pea | 101 (21.4%) |
| **Age (years)** |  |  |  |
| 18–24 | 368 (78.0%) | 35–44 | 10 (2.1%) |
| 25–34 | 71 (15.0%) | 45+ | 23 (4.9%) |
| **Gender** |  |  |  |
| female | 199 (42.2%) | male | 256 (54.2%) |
| non-binary/other | 17 (3.6%) |  |  |
| **Ethnicity** |  |  |  |
| White | 206 (50.9%) | Hispanic/Latino | 46 (11.4%) |
| Asian | 142 (35.1%) | Black/African American | 33 (8.1%) |
| other | 19 (4.7%) |  |  |
| **Dietary Pattern** |  |  |  |
| omnivore | 404 (85.6%) | flexitarian | 28 (5.9%) |
| pescatarian | 24 (5.1%) | vegetarian | 13 (2.8%) |
| vegan | 3 (0.6%) |  |  |
| Characteristic | mean ± SD | | mean ± SD |
| **Meat Attachment** | 3.31 ± 0.91 | **Plant-Based Diet %** | 36.1 ± 22.4 |

### 3.2. Consumer Sensory Attribute Ratings

Table 3 summarizes consumer ratings across the eight sensory attributes, and Figure 2 visualizes these sensory profiles as radar charts. Consumer ratings revealed significant differences across burger formulations for all evaluated attributes. The beef-mushroom burger received the highest ratings for meatiness (77.0 ±19.1), tastiness (69.4 ± 17.1), moistness (61.8 ± 21.7), and softness (58.3 ± 18.1), significantly exceeding all other products (Mann-Whitney U, *p* < 0.05). Meatiness exhibited the largest between-product variation (*H* = 65.76, *p* < 0.001), with ratings ranging from 77.0 for the beef-mushroom burger to 38.6 for the bean burger. The pea burger achieved intermediate meatiness ratings (56.1 ± 20.3), significantly higher than the bean burger but lower than both animal-based burgers. Participants rated the beef-mushroom and turkey burgers similarly for hardness and chewiness, while both products received significantly higher ratings than the pea burger. The bean burger received the lowest ratings for meatiness, fattiness, and moistness, while the pea burger achieved ratings closer to the animal-based burgers for fattiness and moistness than for meatiness and chewiness.

**Table 3:** Consumer sensory attribute ratings. Mean consumer ratings for eight sensory attributes across four burger formulations using 0–100 visual analog scales. Values represent mean ± standard deviation. Kruskal-Wallis H statistics and associated *p*-values quantify overall differences across burger types. Superscript letters indicate results of post-hoc pairwise comparisons; values that share the same letter within a row do not differ significantly (Mann-Whitney U with Bonferroni correction, *p* < 0.05).

| parameter | beef-mushroom | turkey | bean | pea | H | p |
| --- | --- | --- | --- | --- | --- | --- |
| *consumer sensory ratings (mean ± SD)* |  |  |  |  |  |  |
| hard | 55.2 ± 17.8<sup>a</sup> | 53.5 ± 17.1<sup>a</sup> | 50.9 ± 16.8<sup>ab</sup> | 47.3 ± 15.0<sup>b</sup> | 4.85 | 0.003 |
| chewy | 62.4 ± 18.3<sup>a</sup> | 60.6 ± 15.2<sup>a</sup> | 58.8 ± 16.3<sup>ab</sup> | 53.2 ± 16.8<sup>b</sup> | 9.73 | <0.001 |
| moist | 61.8 ± 21.7<sup>a</sup> | 49.2 ± 20.8<sup>bc</sup> | 44.2 ± 18.6<sup>c</sup> | 54.0 ± 17.1<sup>b</sup> | 13.80 | <0.001 |
| fibrous | 50.3 ± 22.0<sup>ab</sup> | 53.5 ± 20.1<sup>a</sup> | 57.3 ± 16.6<sup>a</sup> | 46.0 ± 15.7<sup>b</sup> | 4.58 | 0.004 |
| meaty | 77.0 ± 19.1<sup>a</sup> | 68.0 ± 17.7<sup>b</sup> | 38.6 ± 23.1<sup>d</sup> | 56.1 ± 20.3<sup>c</sup> | 65.76 | <0.001 |
| fatty | 55.3 ± 19.0<sup>a</sup> | 43.5 ± 20.2<sup>b</sup> | 37.0 ± 19.7<sup>b</sup> | 52.9 ± 17.8<sup>a</sup> | 12.42 | <0.001 |
| soft | 58.3 ± 18.1<sup>a</sup> | 47.2 ± 16.7<sup>b</sup> | 49.4 ± 16.6<sup>b</sup> | 48.2 ± 17.1<sup>b</sup> | 12.97 | <0.001 |
| tasty | 69.4 ± 17.1<sup>a</sup> | 58.1 ± 17.3<sup>b</sup> | 60.4 ± 20.3<sup>b</sup> | 56.2 ± 18.9<sup>b</sup> | 17.95 | <0.001 |

**Figure 2:**
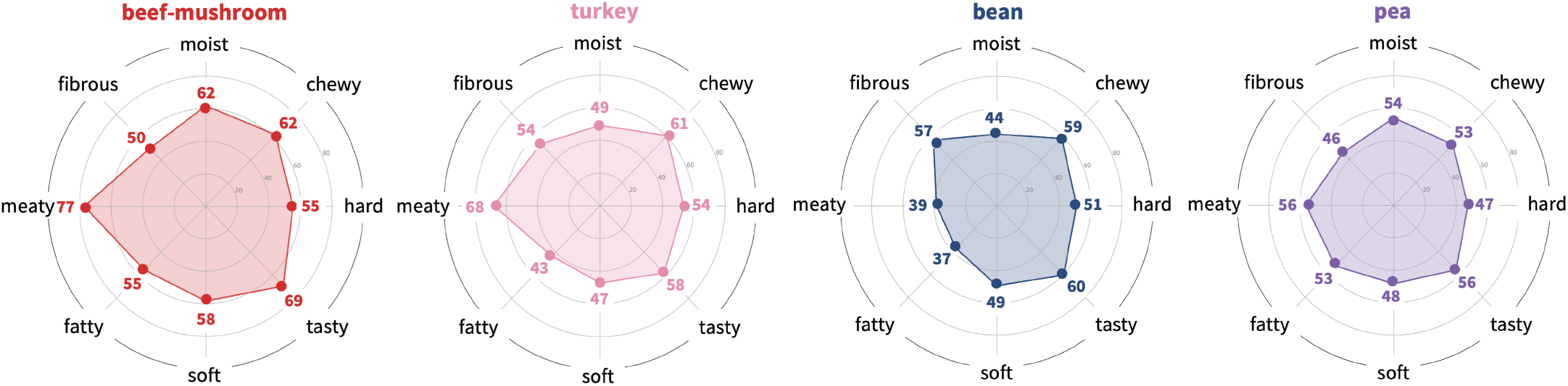
Consumer sensory profiles across burger formulations. Radar charts display mean consumer ratings on 0–100 visual analog scales for eight sensory attributes across four burger types: beef-mushroom, turkey, bean, and pea. The beef-mushroom burger achieved the highest ratings for meatiness, tastiness, moistness, and softness, while the bean burger received the lowest meatiness ratings.

### 3.3. Just-About-Right Penalty Analysis

Figure 3 summarizes the Just-About-Right (JAR) analysis across burger formulations. Moistness emerged as the primary reformulation target for non-beef burgers. The beef-mushroom burger achieved the highest *just about right* percentage for moistness (78%), whereas most participants of the turkey and bean consumers rated moistness as *not enough* (58% and 46%). Penalty analysis quantified the sensory impact of these deviations: insufficient moistness reduced tastiness by 11.2 points for turkey and 13.3 points for bean burgers (*p* < 0.05). The pea burger achieved a more balanced *just about right* moisture profile (63%) but showed substantial penalties for *not enough* savoriness (36%, 21.6-point penalty) and *too much* fibrousness (21%, 18.9-point penalty). In contrast, the beef-mushroom burger showed no actionable deviations that exceeded the 0.50-point threshold on the 7-point overall liking scale.

**Figure 3:**
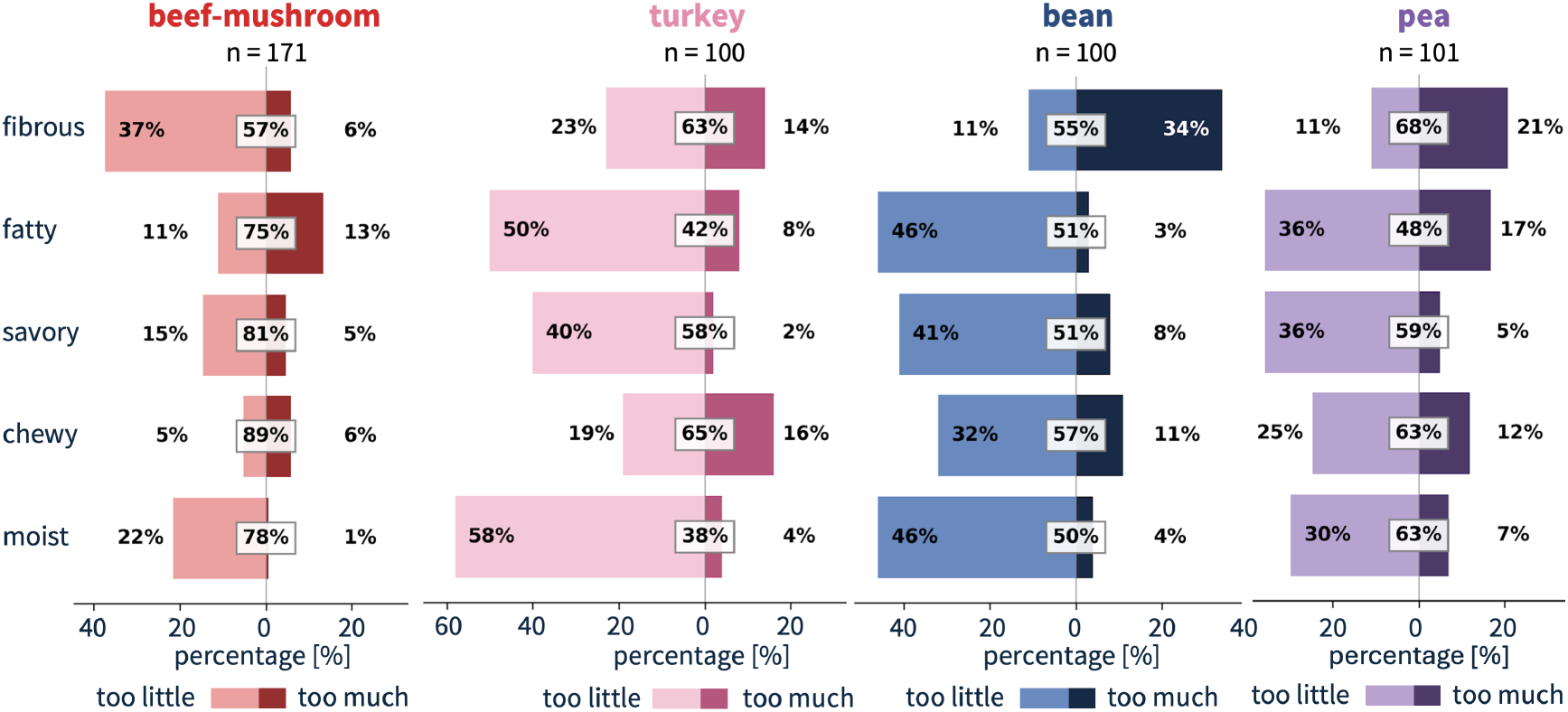
Just-About-Right sensory attribute analysis. Diverging bar charts display the percentage of participants who rated five sensory attributes as *too little* (left), *just about right* (center), or *too much* (right) for the beef-mushroom, turkey, bean, and pea burgers. The beef-mushroom burger achieved the highest fractions of *just about right* ratings in four of the five categories, chewiness (89%), savoriness (81%), moistness (78%), and fattiness (75%). The turkey and bean burgers showed large deficits in perceived moistness (58%) and (46%), and fattiness (50%) and (46%).

### 3.4. Texture Profile Analysis Results

Figure 4 summarizes the texture profile analysis results across the four burgers. Texture profile analysis revealed significant differences across burger types for five of six texture profile analysis parameters. Turkey burgers exhibited the highest stiffness (348.5 ± 65.7 kPa), hardness (8.76 ± 1.65 N), cohesiveness (0.64 ± 0.07), springiness (0.74 ± 0.19), and chewiness (4.26 ± 1.89 N) compared to all other products (Tukey’s HSD, *p* < 0.05). Resilience did not differ significantly across burger types (*p* = 0.12). The bean burger showed the lowest stiffness (97.6 ± 30.0 kPa), hardness (2.45 ± 0.75 N), cohesiveness (0.28 ± 0.03), springiness (0.40 ± 0.09), and chewiness (0.27 ± 0.10 N). The beef-mushroom and pea burgers exhibited intermediate mechanical behavior and did not differ significantly for most texture profile analysis parameters. When grouped by protein source, animal-based burgers (beef-mushroom and turkey) exhibited significantly greater stiffness (*p* < 0.001), hardness (*p* < 0.001), cohesiveness (*p* < 0.001), and chewiness (*p* < 0.001) than plant-based burgers (bean and pea). Springiness (*p* = 0.079) and resilience (*p* = 0.060) did not differ significantly between animal-based and plant-based categories.

**Figure 4:**
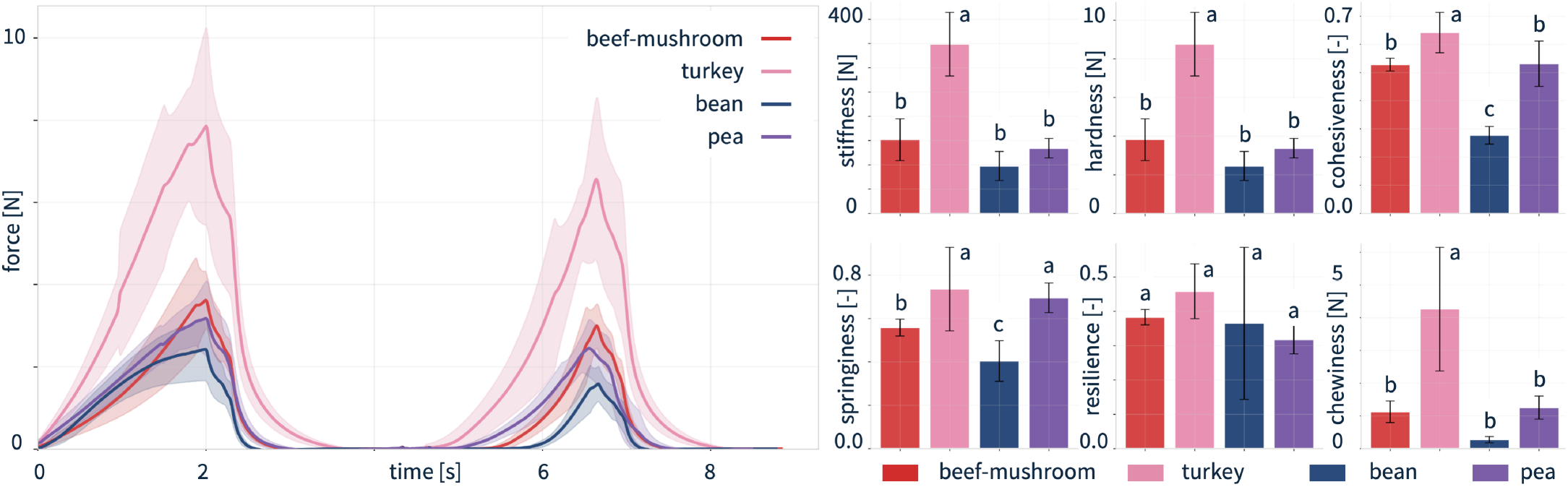
Texture profile analysis results. Representative force-time curves from texture profile analysis double-compression tests for beef-mushroom, turkey, bean, and pea burgers (left). Shaded regions indicate standard deviation (*n* = 10 per product). Turkey burgers exhibited the highest peak forces during both compression cycles, while bean burgers exhibited the lowest mechanical resistance. Comparison of six texture profile analysis parameters across burger types (right). Error bars indicate standard deviation. Letters denote statistically significant differences between groups (Tukey’s HSD, *p* < 0.05).

### 3.5. Sensory Attribute Inter-correlations

Figure 5 illustrates pairwise relationships among sensory attributes within each burger formulation. Across all four products, hardness correlated positively with chewiness, while moistness, fattiness, softness, and meatiness formed a cluster of positively associated attributes. These relationships were particularly pronounced for the beef-mushroom and pea burgers, where consumers who perceived products as more moist, fatty, and meat-like also tended to report higher tastiness. Color coding by overall liking revealed a consistent pattern across burger types: High-liking responses clustered in regions associated with greater meatiness, moistness, fattiness, softness, and tastiness, whereas low-liking responses concentrated at lower values of these attributes. This trend appeared most clearly for the beef-mushroom and pea burgers, and suggests that consumers associated product acceptance with a sensory profile that more closely resembles conventional meat. The bean burger exhibited weaker inter-attribute relationships and a broader dispersion of liking scores, indicating a less cohesive sensory profile. Across all products, hardness and chewiness remained strongly coupled, but these attributes showed weaker associations with overall liking than meatiness, moistness, and tastiness. Exploratory analyses at the burger-product level (*n* = 4) revealed positive relationships between instrumental texture profile analysis parameters and mean consumer sensory ratings. Instrumental cohesiveness showed the strongest association with perceived meatiness (*r* = 0.82) and suggests that products with greater internal structural integrity may appear more meat-like to consumers. Because these analyses included only four products, the resulting correlations should be interpreted as hypothesis-generating rather than confirmatory.

**Figure 5:**
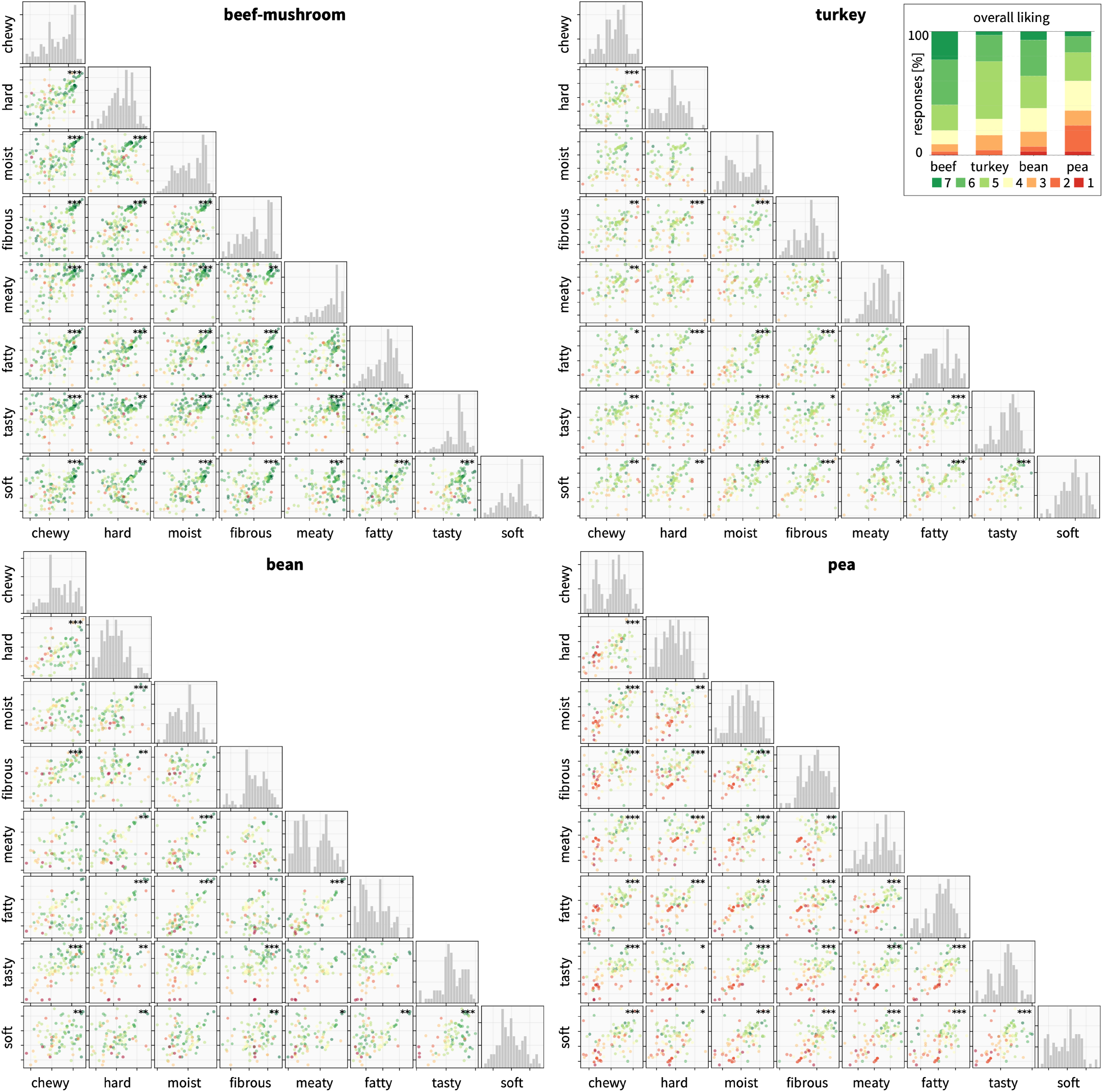
Sensory attribute inter-correlations and overall liking. Lower-triangle scatter matrices show pairwise relationships among eight sensory attributes for beef-mushroom (top left), turkey (top right), bean (bottom left), and pea (bottom right) burgers. Diagonal panels display attribute distributions. Dot color indicates overall liking, from high liking (green) to low liking (red). Across products, higher overall liking clustered with greater meatiness, moistness, fattiness, softness, and tastiness. Significance markers indicate Spearman correlations: *** *p* < 0.001, ** *p* < 0.01, * *p* < 0.05.

### 3.6. Choice Factor Importance

Figure 6 summarizes the importance of factors that influenced burger choice decisions. Flavor emerged as the most important factor overall (71.9 ±16.1), followed closely by texture (67.9 ± 19.3). Health and nutrition ranked third (59.8 ± 24.5), followed by availability (57.1 ± 21.6) and familiarity (51.4 ± 24.7).

**Figure 6:**
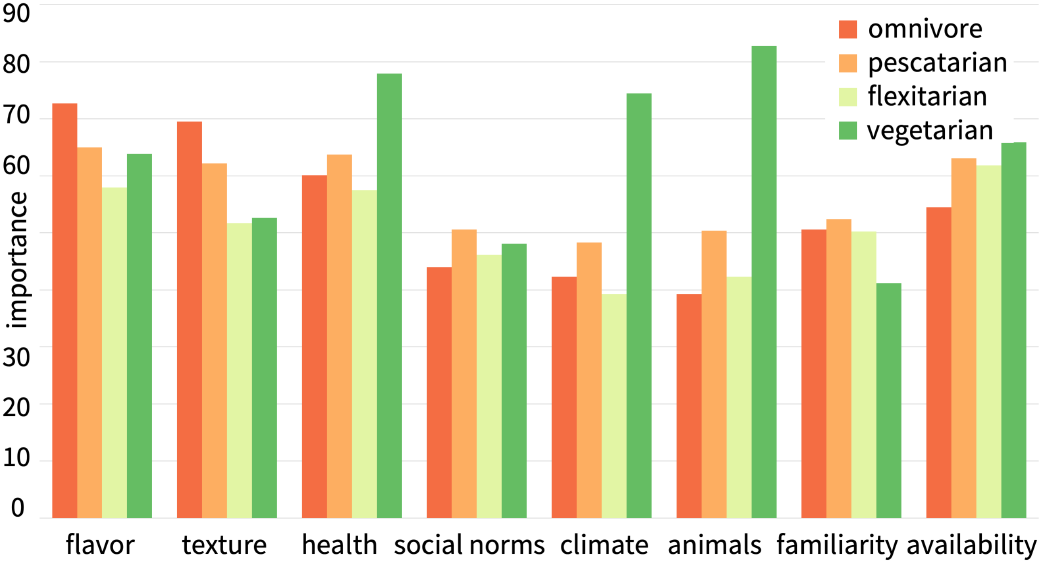
Choice factor importance by dietary pattern. Mean importance ratings (0 −100 scale) for eight factors that influence burger selection across omnivore (*n* = 404), pescatarian (*n* = 24), flexitarian (*n* = 28), and vegetarian (*n* = 13) participants. Flavor and texture ranked highest across all dietary groups, while vegetarians assigned greater importance to health, climate sustainability, and animal welfare than other consumer segments.

Participants assigned lower importance to climate sustainability (44.7 ± 21.9), social norms (44.6 ± 23.1), and animal welfare (43.0 ± 23.6). Dietary patterns influenced these priorities. Omnivores placed the greatest emphasis on flavor and texture, whereas vegetarians assigned substantially greater importance to health, climate sustainability, and animal welfare.

### 3.7. Demographic Analyses

Meat Attachment Questionnaire (MAQ) scores differed significantly across dietary patterns (Kruskal-Wallis *H* = 21.43, *p* < 0.001). Omnivores reported the strongest attachment to meat (3.41 ± 0.91), followed by flexitarians (3.05 ± 0.85), pescatarians (2.68 ± 0.76), vegans (2.40 ± 1.22) and vegetarians (2.04 ± 0.90). MAQ scores also differed significantly across burger choices (*H* = 25.80, *p* < 0.001). Participants who selected the beef-mushroom burger reported higher meat attachment (3.58 ± 0.90) than participants who selected pea (3.03 ± 0.88) or bean (3.11 ± 0.90) burgers. These findings suggest that individual attitudes toward meat influenced both dietary habits and burger selection.

## 4. Discussion

This study combined real-world consumer sensory evaluation with instrumental texture profile analysis to identify texture drivers of burger acceptance across blended, animal-based, and plant-based formulations. Three findings emerged: First, the beef-mushroom burger achieved the strongest consumer response, with the highest ratings for meatiness, tastiness, moistness, and softness. Second, instrumental texture alone did not fully explain consumer acceptance; the pea burger showed mechanical properties similar to the beef-mushroom burger but received lower sensory ratings. Third, moistness and cohesiveness emerged as key formulation targets that link consumer perception to measurable physical properties.

### Texture shapes acceptance, but mechanics alone do not determine liking

Texture remains a fundamental challenge in the development of meat alternatives and blended products [26]. The hierarchical structure of animal muscle–with muscle fibers, connective tissue, fat, and water–creates a sensory experience that plant-based materials often struggle to reproduce [27]. Consistent with previous consumer studies, our results confirm that textural attributes such as juiciness, meatiness, and fibrousness strongly influence acceptance of meat analogues [29]. Our texture profile analysis revealed substantial differences in mechanical behavior across burger types, with turkey burgers exhibiting the highest stiffness, hardness, and chewiness. By contrast, the pea burger showed mechanical properties that more closely resembled the beef-mushroom burger than the bean burger. This convergence suggests that modern protein structuring technologies, including extrusion of pea protein, can successfully reproduce several mechanical features of animal-derived products [4]. However, the lower cohesiveness of the bean burger indicates a more brittle and less integrated matrix that may contribute to crumbling during mastication. Despite these mechanical similarities, the pea burger did not match the beef-mushroom burger in perceived meatiness or overall sensory performance. This discrepancy highlights an important distinction between mechanical texture and consumer perception: *Mechanical parity does not guarantee sensory parity*. Flavor release, fat perception, moisture distribution, and oral processing likely contribute alongside mechanics to shape the overall eating experience.

### Cohesiveness links structure to perceived meatiness

Among the instrumental texture parameters, cohesiveness showed the strongest association with perceived meatiness (*r* = 0.82). This relationship suggests that internal structural integrity contributes to a meat-like eating experience. *Cohesiveness may represent a mechanical signature of perceived meatiness*. Texture profile analysis was originally developed to quantify sensory-relevant properties such as cohesiveness, springiness, and chewiness [25]. Recent work on plant-based and animal proteins has highlighted cohesiveness as an important descriptor of structural integrity and eating quality [19]. In agreement with previous studies, the bean burger had the lowest cohesiveness, consistent with a more brittle and crumbly matrix, while the beef-mushroom and pea burgers achieved substantially higher values [7]. Similar structure-function relationships have emerged in studies of fibrous plant-protein architectures [30]. Together, these findings identify cohesiveness as a useful mechanical target for formulation design and suggest that products with insufficient binding may struggle to deliver the structural continuity that consumers associate with meat.

### Moisture retention is the clearest reformulation target

The Just-About-Right analysis identified moistness as the most actionable sensory limitation for non-beef burgers. More than half of turkey consumers rated the product as not moist enough, and nearly half of bean consumers reported the same deficit. In agreement with previous findings, these deviations carried substantial penalties for tastiness [31]. *Consumers strongly penalize products that lack moisture*. In contrast, the beef-mushroom burger achieved the highest moistness rating and the highest proportion of just-about-right moistness responses. Moisture retention and juiciness consistently rank among the strongest predictors of consumer acceptance in meat alternatives [32]. Recent sensory studies of plant-based proteins similarly identify moistness as a major driver of overall liking [33]. Our findings reinforce the importance of water-binding capacity, fat structuring, and controlled moisture release during mastication.

### Blended burgers offer a practical transition strategy

By combining grass-fed beef with mushrooms, the beef-mushroom burger represents a blended approach that may appeal to flexitarian consumers who seek to reduce meat consumption without fully abandoning animal products [34]. It outperformed the other formulations across the most important sensory dimensions, including meatiness (77.0/100), tastiness (69.4/100), moistness (61.8/100). Its strong consumer performance suggests that blended products can reduce meat content while preserving sensory cues that consumers value. Previous studies of blended meat-mushroom products report consumer acceptance levels comparable to conventional meat products despite lower animal protein content [14]. Recent analyses identify blended products as a promising pathway toward dietary transition because they maintain familiar sensory characteristics while improving sustainability metrics [17]. Importantly, the beef-mushroom burger maintained cohesiveness and meatiness despite partial substitution with mushrooms [15]. *Blended products may offer the most practical pathway to reduce meat consumption without sacrificing consumer acceptance*. This finding supports blended products as a pragmatic strategy for incremental meat reduction, especially among consumers who remain attached to conventional meat.

### Consumer preferences reflect both product properties and self-selection

The dining hall design increased ecological validity because participants selected burgers under normal meal conditions. At the same time, this design introduced self-selection: Participants who chose the beef-mushroom burger reported higher meat attachment (3.58) than those who chose pea (3.03) or bean (3.11) burgers. Importantly, omnivores reported significantly higher meat attachment (3.41) than flexitarians (3.05) and vegetarians (2.04), indicating that psychological attachment to meat represents a meaningful barrier that product developers must address through sensory optimization. This result agrees with previous studies in which stronger meat attachment influenced both food choice and willingness to reduce meat consumption [23]. Dietary identity and motivational factors also shape attitudes toward meat alternatives [35]. *Consumer acceptance reflects both product quality and consumer identity*. These differences suggest that product ratings partly reflect pre-existing consumer preferences, not only intrinsic product properties. Future within-subjects studies could separate product effects from consumer selection effects.

### Implications for product development

These findings suggest three practical design principles: First, product developers should treat cohesiveness as a key mechanical target because it tracks perceived meatiness and reflects internal structural integrity. Second, developers should prioritize moisture retention and moisture release because consumers strongly penalize dry products. Third, developers should not rely on mechanical matching alone. The pea burger resembled the beef-mushroom burger in several texture profile analysis parameters, but did not achieve comparable consumer acceptance. *Successful meat alternatives must optimize sensory perception, not mechanical properties alone*. Emerging frameworks increasingly view food as an engineered material whose performance depends on the interplay of structure, mechanics, and sensory perception [18]. Recent studies of meat analogues similarly emphasize the need to combine mechanical targets with flavor and juiciness optimization [19].

### Limitations and scope

This study has several limitations: First, the dining hall setting increased ecological validity but introduced variation in food holding time, serving conditions, toppings, and eating context. Second, the Stanford community may not represent the broader U.S. population. Third, each participant evaluated only one burger, which limits causal comparisons across formulations. Fourth, texture profile analysis relied on excised cylindrical samples and therefore did not fully capture the experience of biting through an intact burger. Fifth, correlations between texture profile analysis parameters and sensory ratings were based on only four products and should be interpreted as hypothesis-generating.

### Toward sustainable protein transitions

Future studies should evaluate burgers in within-subjects sensory designs, test larger product panels, and quantify how toppings, condiments, buns, and serving temperature influence sensory perception.

For blended burgers, systematic variation of mushroom-to-beef ratio could identify the threshold at which sensory and mechanical properties diverge from conventional beef. Reducing meat consumption remains an important opportunity to improve both environmental sustainability and public health. Based on the present results, future formulations should target high moisture retention and cohesiveness while preserving the sensory attributes that consumers associate with meat. Ultimately, successful protein transitions will depend not only on reducing environmental impact, but also on delivering products that people genuinely enjoy eating.

## 5. Conclusion

This study combined consumer sensory evaluation with instrumental texture profile analysis to identify the factors that drive burger acceptance across blended, animal-based, and plant-based formulations. The beef-mushroom burger achieved the strongest consumer response with the highest ratings for meatiness, tastiness, and moistness. Although the pea burger matched the beef-mushroom burger in several mechanical properties, it did not achieve comparable sensory performance which indicates that texture parity alone does not ensure consumer acceptance. Across formulations, moistness emerged as the primary sensory limitation, while cohesiveness showed the strongest relationship with perceived meatiness and internal structural integrity. These findings suggest that successful burger reformulation requires more than mechanical matching; products must deliver the moisture, flavor, and eating experience that consumers associate with meat. Blended formulations appear particularly effective because they preserve these sensory qualities while reducing animal protein content. Blended products offer a practical pathway to reduce meat consumption because they preserve the sensory experience of meat and provide a realistic bridge toward more sustainable diets.

## Data availability

De-identified sensory data, mechanical texture analysis results, and data processing scripts are publicly available at https://github.com/LivingMatterLab/AI4Food.

## Acknowledgments

The authors thank Andrew Mayne, Senior Associate Director of Culinary Event Strategy & Plant Forward Experiences at Stanford University and Stanford Residential & Dining Enterprises for facilitating data collection. This research was supported by the Stanford Bio-X Snack Grant 2025, the Stanford SDSS Accelerator Grant 2025, the NSF CMMI Award 2320933, and the ERC Advanced Grant 101141626.

